## Supplementary Material for "Multi-study fMRI outlooks on subcortical BOLD responses in the stop-signal paradigm"

### Appendix

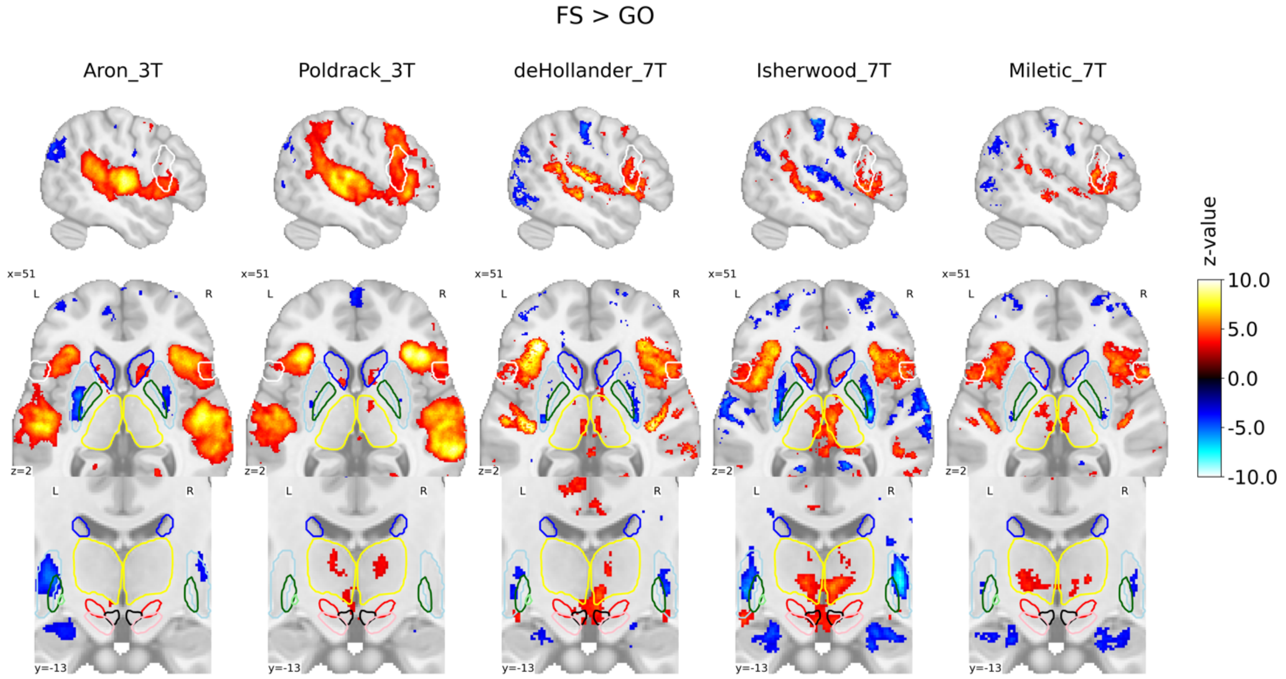


**Supplementary Figure 1.** Group-level SPMs of the FS > GO contrast of the SST for each dataset. Activation colours indicate FDR thresholded (q < .05) z-values. Sagittal (top), axial (middle) and a zoomed in coronal (bottom) view are shown. Coloured contour lines indicate regions of interest (IFG in white, M1 in grey, preSMA in orange, Caudate in dark blue, Putamen in light blue, GPe in dark green, GPi in light green, SN in pink, STN in red, thalamus in yellow, and VTA in black). The background template and coordinates are in MNI2009c (1mm); slices are drawn through x = 51 (top), y = -13 (bottom), and z = 2 (middle).

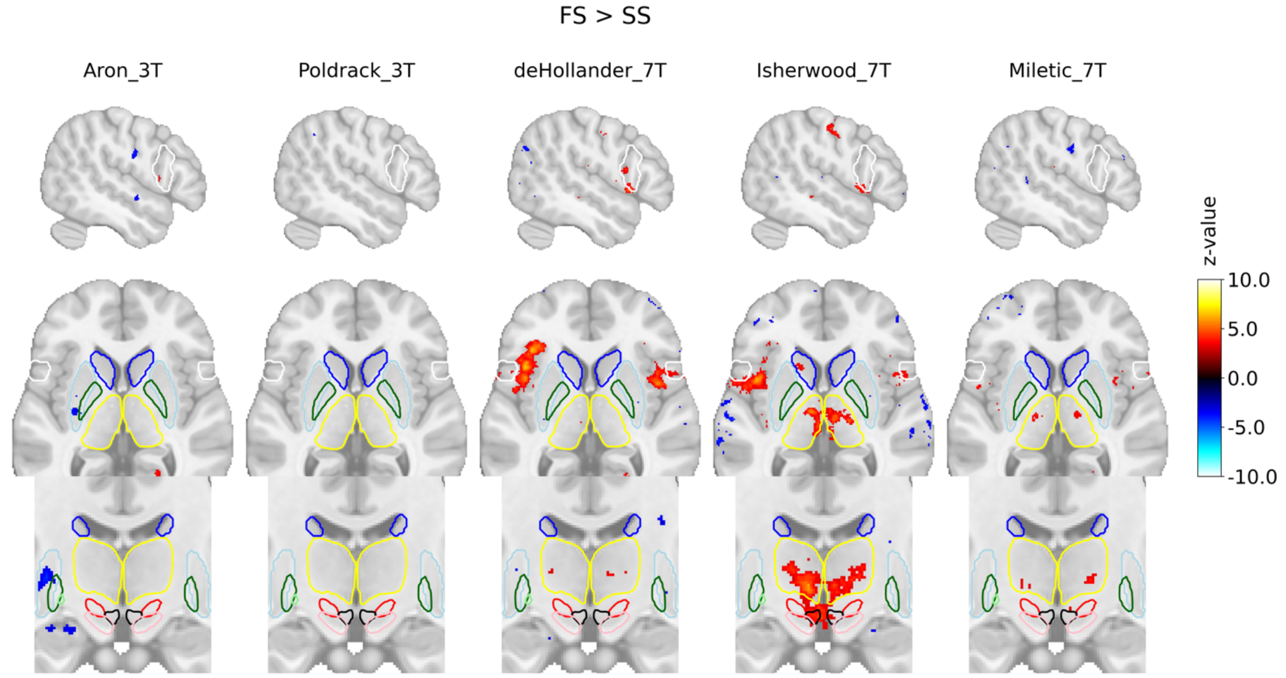


**Supplementary Figure 2.** Group-level SPMs of the FS > SS contrast of the SST for each dataset. Activation colours indicate FDR thresholded (q < .05) z-values. Sagittal (top), axial (middle) and a zoomed in coronal (bottom) view are shown. Coloured contour lines indicate regions of interest (IFG in white, M1 in grey, preSMA in orange, Caudate in dark blue, Putamen in light blue, GPe in dark green, GPi in light green, SN in pink, STN in red, thalamus in yellow, and VTA in black). The background template and coordinates are in MNI2009c (1mm); slices are drawn through x = 51 (top), y = -13 (bottom), and z = 2 (middle).

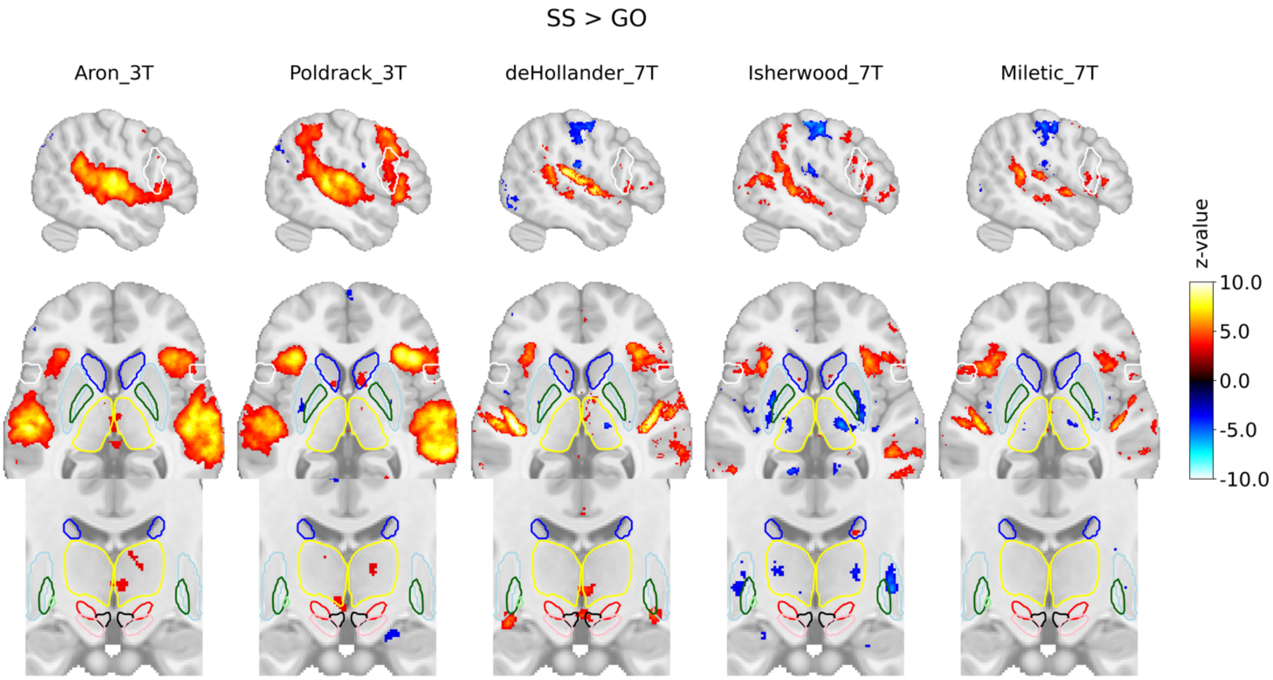


**Supplementary Figure 3.** Group-level SPMs of the SS > GO contrast of the SST for each dataset. Activation colours indicate FDR thresholded ($q < .05$) z-values. Sagittal (top), axial (middle) and a zoomed in coronal (bottom) view are shown. Coloured contour lines indicate regions of interest (IFG in white, M1 in grey, preSMA in orange, Caudate in dark blue, Putamen in light blue, GPe in dark green, GPi in light green, SN in pink, STN in red, thalamus in yellow, and VTA in black). The background template and coordinates are in MNI2009c (1mm); slices are drawn through x = 51 (top), y = -13 (bottom), and z = 2 (middle).

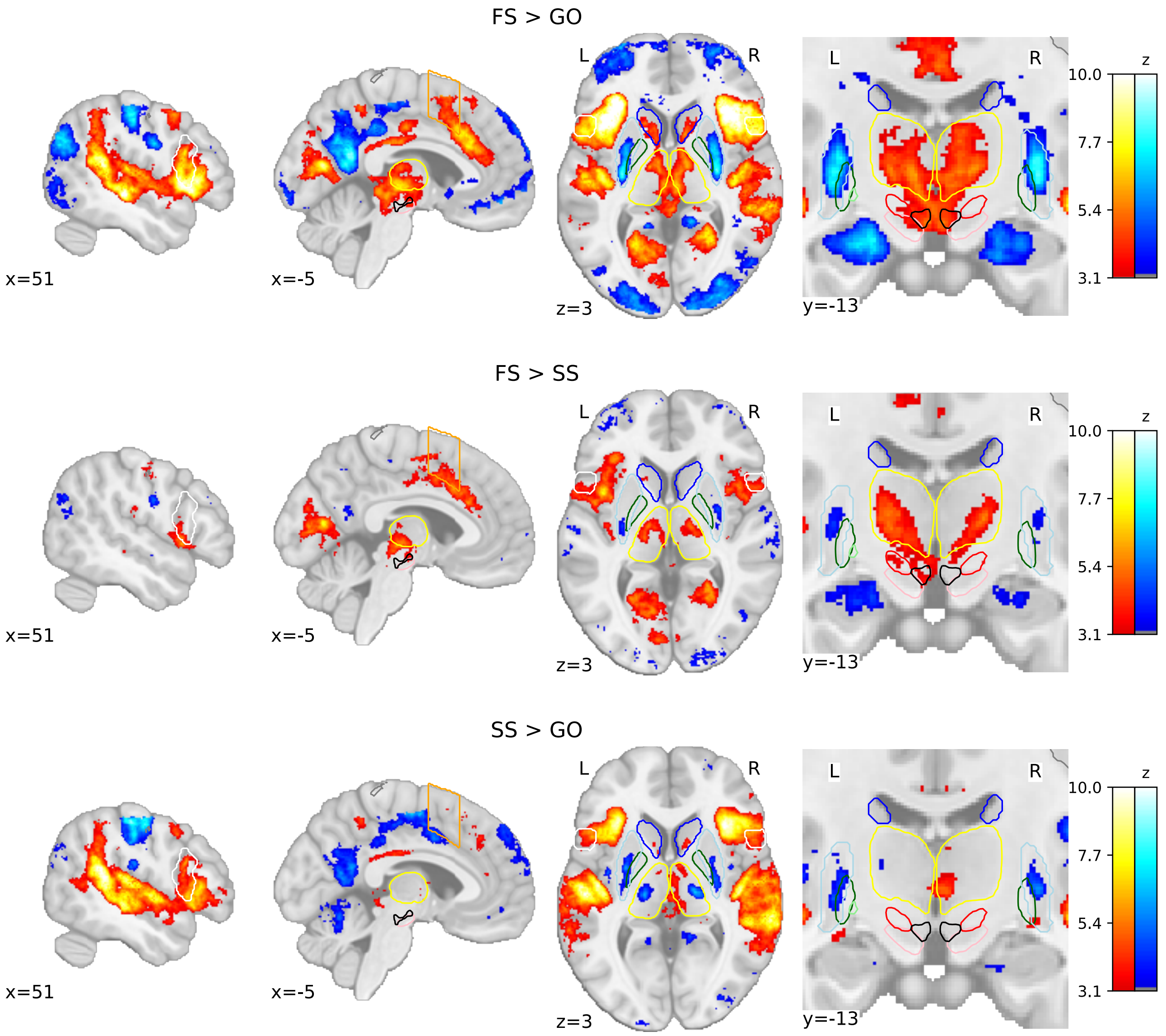


**Supplementary Figure 4.** Group-level SPMs of the three main contrasts of the SST, where SS and FS trials were time-locked to the presentation of the stop signal. Activation colours indicate FDR thresholded ($q < .05$) z-values. Two sagittal, one axial, and one zoomed in coronal view are shown. Coloured contour lines indicate regions of interest (IFG in white, M1 in grey, preSMA in orange, Caudate in dark blue, Putamen in light blue, GPe in dark green, GPi in light green, SN in pink, STN in red, thalamus in yellow, and VTA in black). The background template and coordinates are in MNI2009c (1mm). FS, failed stop; SS, successful stop.

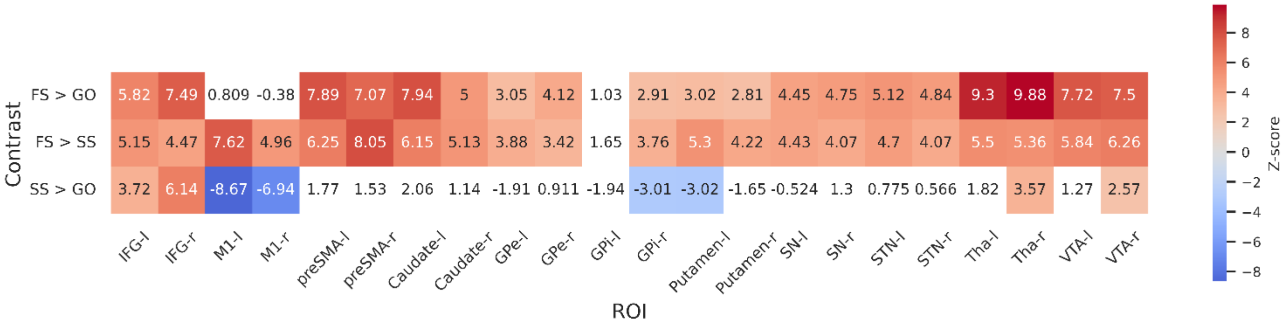


**Supplementary Figure 5.** Group-level z-scores from the ROI-wise GLM analysis of included datasets, where SS and FS trials were time-locked to the presentation of the stop signal. Thresholds are set using FDR correction ($q < .05$), varying between contrasts. The thresholds for each contrast are as follows: 3.01 for FS > GO, 2.26 for FS > SS and 3.1 for SS > GO. Regions that do not reach significance are coloured white. Left and right hemispheres are shown separately, denoted by '-l' or '-r', respectively. IFG, inferior frontal gyrus; M1, primary motor cortex; preSMA, pre-supplementary motor area; GPe, globus pallidus externa; GPi, globus pallidus interna; SN, substantia nigra; STN, subthalamic nucleus; Tha, thalamus; VTA, ventral tegmental area.

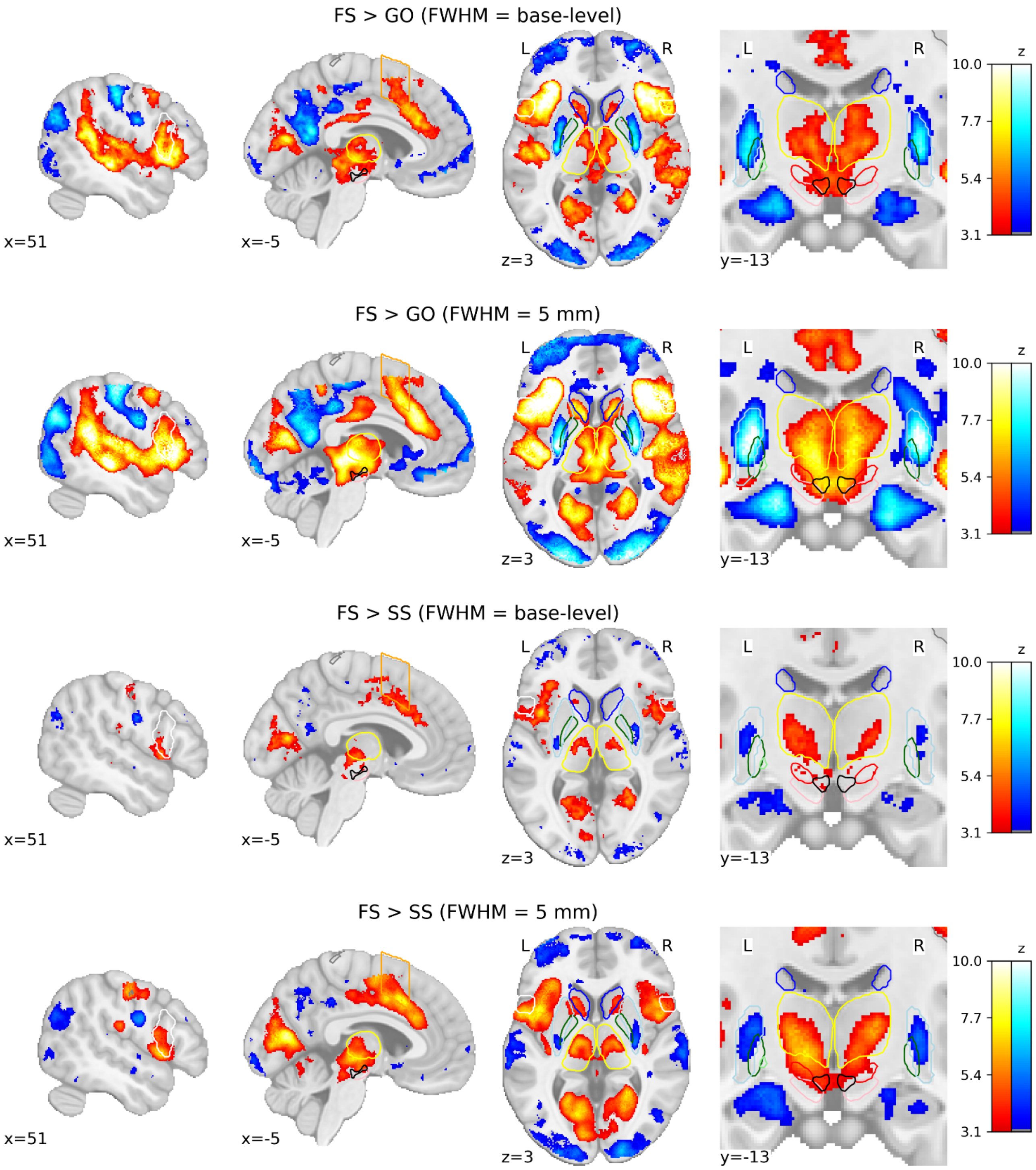


**Supplementary Figure 6.** Group-level SPMs of the FS > GO and FS > SS contrasts using different smoothing kernels. Activation colours indicate FDR thresholded (q < .05) z-values. Two sagittal, one axial, and one zoomed in coronal view are shown. Coloured contour lines indicate regions of interest (IFG in white, M1 in grey, preSMA in orange, Caudate in dark blue, Putamen in light blue, GPe in dark green, GPi in light green, SN in pink, STN in red, thalamus in yellow, and VTA in black). The background template and coordinates are in MNI2009c (1mm). FS, failed stop; SS, successful stop.

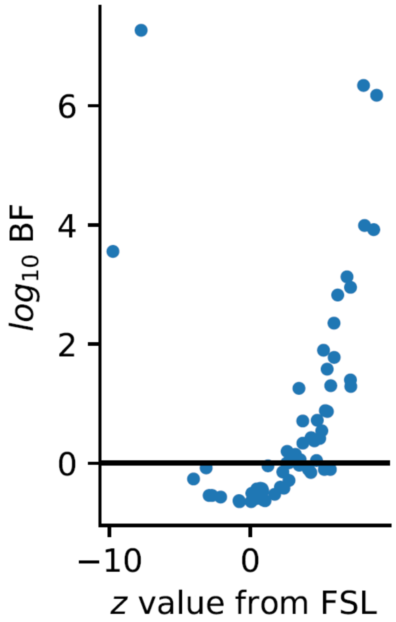


**Supplementary Figure 7.** A comparison of the BFs and the frequentist *z*-scores from FSL. The 0 point on each axis represents no evidence for an effect. Large absolute z-values are expected to also yield a high log BF, hence the inverted U-shape shows good correspondence between the two estimates.
